# Cone-like rhodopsin expressed in the all cone retina of the colubrid pine snake as a potential adaptation to diurnality

**DOI:** 10.1101/100792

**Authors:** Nihar Bhattacharyya, Benedict Darren, Ryan K. Schott, Vincent Tropepe, Belinda S.W. Chang

**Affiliations:** Department of Cell & Systems Biology, University of Toronto, Toronto, Ontario, Canada; Department of Ecology & Evolutionary Biology, University of Toronto, Toronto, Ontario, Canada; Department of Ophthalmology & Vision Sciences, University of Toronto, Toronto ON, Canada M5T 3A9; Centre for the Analysis of Genome Evolution and Function, University of Toronto, Toronto, Ontario, Canada

**Keywords:** rod and cone photoreceptors, photoreceptor transmutation, rhodopsin, visual pigments, visual evolution, reptile vision

## Abstract

Colubridae is the largest and most diverse family of snakes, with visual systems that reflect this diversity, encompassing a variety of retinal photoreceptor organizations. The transmutation theory proposed by Walls postulates that photoreceptors could evolutionarily transition between cell types in squamates, but few studies have tested this theory. Recently, evidence for transmutation and rod-like machinery in an all cone retina has been identified in a diurnal garter snake (*Thamnophis*), and it appears that the rhodopsin gene at least may be widespread among colubrid snakes. However, functional evidence supporting transmutation beyond the existence of the rhodopsin gene remains rare. We examined the all cone retina of another diurnal colubrid, *Pituophis melanoleucus*, distantly related to *Thamnophis*. We found that *P. melanoleucus* expresses two cone opsins (SWS1, LWS) and rhodopsin (RH1) within the eye. Immunohistochemistry localized rhodopsin to the outer segment of photoreceptors in the all-cone retina of the snake and all opsin genes produced functional visual pigments when expressed *in vitro*. Consistent with other studies, we found that *P. melanoleucus* rhodopsin is extremely blue-shifted. Surprisingly, *P. melanoleucus* rhodopsin reacted with hydroxylamine, a typical cone opsin characteristic. These results support the idea that the rhodopsin-containing photoreceptors of *P. melanoleucus* are the products of evolutionary transmutation from rod ancestors, and suggests that this phenomenon may be widespread in colubrid snakes. We hypothesize that transmutation may be an adaptation for diurnal, brighter-light vision, which could result in increased spectral sensitivity and chromatic discrimination with the potential for colour vision.

**Summary Statement:** The all cone retina of the colubrid snake, *Pituophis melanoleucus* contains a blue-shifted rhodopsin with cone opsin-like properties, which may have been adaptive in diurnal snakes.

## Introduction

Reptiles are known for their impressive array of visual adaptations and retinal organizations, which reflect distinct ecologies and evolutionary histories (Walls, 1942). The family Colubridae is the most speciose family of snakes and encompasses a diverse range of lifestyles and ecologies. Colubrid snakes have recently emerged as a compelling group in which to study visual system evolution and adaptation (Schott et al., 2016; Simões et al., 2015; Simões et al., 2016).

In the vertebrate retina, photoreceptor cells can be divided into two types based on their overall structure and function: cones, which are active in bright light and contain cone visual pigments (SWS1, SWS2, RH2, LWS) in a tapered outer segment, and rods, which function in dim light and contain rhodopsin (RH1) in a longer, more cylindrical outer segment (Bowmaker, 2008; Lamb, 2013; Walls, 1942). Reptilian retinas are unique in having multiple retinal configurations among closely related species including all-rod (Kojima et al., 1992), rod and cone (Sillman et al., 2001), and all-cone (Sillman et al., 1997). In 1942, physiologist Gordon Walls outlined his theory of transmutation to explain the evolutionary transformation of photoreceptors from one type to another (Walls, 1942). This phenomenon has since been investigated in nocturnal geckos, where cone opsins are expressed in an all-rod retina in order to compensate for the evolutionary loss of RH1 in a hypothesized diurnal, all-cone ancestor (Kojima et al., 1992; Taniguchi et al., 1999). While the nocturnal henophedian snakes, such as boas and pythons, are known to have duplex retinas expressing RH1, LWS and SWS1 in canonical photoreceptors (Davies et al., 2009), the more derived diurnal colubrid snakes have been primarily shown to posses simplex retinas comprising of all cone photoreceptors, with the fate of the rod photoreceptor unknown (Caprette, 2005; Walls, 1942). Early studies of the colubrid visual system found a green-sensitive visual pigment in addition to a red and a blue pigment (Sillman et al., 1997) in the simplex retina, but were unable to distinguish between a spectrally shifted rhodopsin in a transmuted rod or a potentially resurrected RH2 cone opsin (Cortesi et al., 2015). More recently, a study from our group identified a functional blue-shifted RH1 pigment in the all-cone retina of the ribbon snake (*Thamnophis proximus*), and proposed that this resulted from a rod to cone evolutionary transmutation in colubrid snakes that may have allowed for enhanced spectral discrimination and even trichromatic colour vision (Schott et al., 2016). A recent study that sequenced the opsins of several other colubrid snake species discovered the widespread presence of full-length rhodopsin genes in species with supposed simplex retinas that were previously presumed to have lost rod/rhodopsins (Simões et al., 2016). However, detailed characterizations of colubrid snake opsins and photoreceptors in the context of the theory of evolutionary transmutation still remain rare.

To further test the hypothesis of widespread transmutation in colubrid snakes, and its potential functional consequences, we examined the visual system of the Northern Pine Snake (*Pituophis melanoleucus*), a diurnal colubrid snake distantly related to *T. proximus. Pituophis melanoleucus* inhabits the eastern half of the United States and Canada (Stull, 1940) and spends relatively short intervals on the surface during the day to forage for prey such as small mammals and birds, and to create new burrows (Diller and Wallace, 1996; Himes, 2001). While *P. melanoleucus* has been found to possess an all-cone retina (Caprette, 2005), similar to previous diurnal colubrids snakes studied (Schott et al., 2016; Sillman et al., 1997), unlike other strongly diurnal colubrids such as the garter snake, *P. melanoleucus* is more secretive and is thought to spend a considerable amount of time burrowing (Gerald et al., 2006).

In this study, we investigate whether there is evidence of photoreceptor transmutation from rods into cones in the all-cone retina of *P. melanoleucus* via functional characterization, cellular localization, and molecular evolutionary analyses of its visual pigment (opsin) genes. We isolated three opsins genes from *P. melanoleucus*: SWS1, LWS and RH1. Immunohistochemistry of the retina localized rhodopsin (RH1) protein and rod transducin to the inner and outer segments of a small subset of photoreceptors, suggesting that *P. melanoleucus* supports the theory of rod-to-cone transmutation in diurnal colubrids. All three opsins were successfully expressed *in vitro* and displayed properties characteristic of fully functional visual pigments. Additionally, spectroscopic assays revealed that *P. melanoleucus* rhodopsin is sensitive to hydroxylamine, which is more typical of cone opsins and is suggestive of more cone-like functional properties. This study provides further evidence for a fascinating evolutionary transformation in the retinas of colubrid snakes, with implications for reptiles in general.

## Materials and Methods

### Animals

A Northern pine snake (*Pituophis melanoleucus melanoleucus*, adult) specimen and mice (*Mus musculus*, adult, CD1) were obtained from a licensed source as commissioned by the University Animal Care Committee (UACC). The specimen was sacrificed using an approved euthanasia protocol. The eyes were enucleated and preserved in RNAlater or 4% paraformaldehyde.

### Total RNA extraction and cDNA synthesis

The dissected whole eye was homogenized with TRIzol, and total RNA was isolated using a phenol/chloroform extraction and ethanol precipitation. The first strand of complementary DNA (cDNA) was synthesized using SuperScript III Reverse Transcriptase (Invitrogen, Waltham, MA, USA) from RNA samples primed with a 3’ oligo-dT and a 5’ SMART primer, following the protocol outlined by the SMART cDNA Library Construction Kit (BD Biosciences, Franklin Lakes, NJ, USA). The second strand complement was synthesized by long-distance PCR following the same protocol.

Visual pigment genes were isolated using a degenerate PCR strategy. Degenerate primers based on an alignment of reptilian visual pigment sequences were used in attempts to amplify partial sequences of the LWS, SWS1 and RH1 opsin genes with a heminested strategy. GenomeWalker (Clontech, Mountain View, CA, USA) was additionally used to obtain full-length sequences (Supplementary Table S1). Extracted PCR products were ligated into the pJET1.2 blunt plasmid vector.

### Phylogenetic analysis

A representative set of vertebrate visual opsin sequences was obtained from Genbank. These sequences were combined with the three opsins genes sequenced from the pine snake and aligned using MUSCLE (Edgar, 2004). The poorly aligned 5ʹ and 3ʹ ends of the sequence were manually trimmed. Species list and accession numbers for all sequences used in the study are provided in Supplementary Table S2. In order to confirm the identities of the opsin genes from the pine snake, a gene tree was estimated using the resulting alignment in MrBayes 3 (Ronquist and Huelsenbeck, 2003) using reversible jump MCMC with a gamma rate parameter (nst=mixed, rates=gamma), which explores the parameter space for the nucleotide model and the phylogenetic tree simultaneously. The analyses were run for five million generations with a 25% burn-in. Convergence was confirmed by checking that the standard deviations of split frequencies approached zero and that there was no obvious trend in the log likelihood plot.

### Protein expression

Full-length opsins sequences (RH1, SWS1, and LWS) were amplified from pJET1.2 vector using primers that added the *BamHI* and EcoRI restriction sites to its 5ʹ and 3ʹ ends, respectively, and inserted into the p1D4-hrGFP II expression vector (Morrow and Chang, 2010). Expression vectors containing *P. melanoleucus* cone opsins and rhodopsin genes were transiently transfected into cultured HEK293T cells (ATCC CRL-11268) using Lipofectamine 2000 (Invitrogen, Waltham, MA, USA; 8 μg of DNA per 10-cm plate) and harvested after 48 h. Visual pigments were regenerated with 11-cis retinal, generously provided by Dr. Rosalie Crouch (Medical University of South Carolina), solubilized in 1% dodecylmaltoside, and purified with the 1D4 monoclonal antibody (University of British Columbia #95-062, Lot #1017; Molday and MacKenzie, 1983) as previously described (Morrow and Chang, 2015; Morrow and Chang, 2010; Morrow et al., 2011). RH1 and SWS1 pigments were purified in sodium phosphate buffers and LWS was purified in HEPES buffers containing glycerol (as described in van Hazel et al. (2013)). The ultraviolet-visible absorption spectra of purified visual pigments were recorded using a Cary 4000 double beam spectrophotometer (Agilent, Santa Clara, CA, USA). Dark-light difference spectra were calculated by subtracting light-bleached absorbance spectra from respective dark spectra. Pigments were photoexcited with light from a fiber optic lamp (Dolan-Jenner, Boxborough, MA, USA) for 60 s at 25°C. Absorbance spectra for acid bleach and hydroxylamine assays were measured following incubation in hydrochloric acid (100mM) and hydroxylamine (NH_2_OH, 50mM), respectively. To estimate λ_max_, the dark absorbance spectra were baseline corrected and fit to Govardovskii templates for A1 visual pigments (Govardovskii et al., 2000).

### Immunohistochemistry

Fixation of pine snake eyes was conducted as previously described (Schott et al., 2016). Briefly, after enucleating *P. melanoleucus* eyes in the light, they were rinsed in PBS (0.8% NaCl, 0.02% KCl, 0.144% NaHPO4, and 0.024% KH2PO4, pH 7.4), fixed overnight at 4°C in 4% paraformaldehyde, infiltrated with increasing concentrations of sucrose (5%, 13%, 18%, 22%, 30%) in PBS, and embedded in a 2:1 solution of 30% sucrose and O.C.T compound (Tissue-Tek, Burlington, NC, USA) at −20°. The eyes were cryosectioned transversely at −25°C in 20 μm sections using a Leica CM3050 (Wetzlar, Germany) cryostat, placed onto positively charged microscope slides, and stored at −80°C until use.

Slides were first rehydrated in PBS and then air-dried to ensure adhesion. Sections were rinsed three times in PBS with 0.1% Tween-20 (PBT) and then incubated in 4% paraformaldehyde PBS for 20 minutes. After rinsing in PBT and PDT (PBT with 0.1% DMSO), the slides were incubated in a humidity chamber with blocking solution (1% BSA in PDT with 2% normal goat serum) for one hour, incubated with primary antibody diluted in blocking solution overnight at 4° in a humidity chamber. Antibodies used were the K20 antibody (Santa Cruz Biotechnology, Santa Cruz, CA, USA, sc-389, lot#:C1909, dilution: 1:500) and RET-P1 anti-rhodopsin antibody (Sigma-Aldrich, St. Louis, MO, USA, O-4886, lot#: 19H4839, dilution: 1:200).

After extensive rinsing and soaking in PDT (3 times for 15 minutes), secondary antibody was added to the samples and incubated at 37° for one hour in a humidity chamber. Secondary antibodies used for the fluorescent staining were the AlexaFluor-488 anti-rabbit antibody (Life Technologies, Waltham, MA, USA, A11034, lot#: 1298480, dilution: 1:1000) and the Cy-3 anti-mouse antibody (Jackson ImmunoResearch, West Grove, PA, USA, 115-165-003, dilution: 1:800). After rinsing with PBS, followed by PDT, sections were stained with 10 μg/mL Hoescht for 10 minutes at room temperature. The sections were then rinsed in PBS and PDT and mounted with ProLong^®^ Gold antifade reagent (Life technologies, Waltham, MA, USA) and coverslipped. Sections were visualized via a Leica SP-8 confocal laser microscope (Wetzlar, Germany).

## Results

### Full-length RH1, SWS1 and LWS opsin sequences found in Pituophis melanoleucus cDNA

To determine the identities of the visual pigments in *P. melanoleucus*, eye cDNA and gDNA was screened for opsin genes. Three full-length opsins were amplified, sequenced, and analyzed phylogenetically with a set of representative vertebrate visual opsins (Table S1) using Bayesian inference (MrBayes 3.0) (Ronquist and Huelsenbeck, 2003). This analysis confirmed the identity of the three opsin genes as RH1, LWS, and SWS1 (Fig. S1-S3).

All three opsin genes sequence contained the critical amino acid residues required for proper structure and function of a prototypical opsin including K296, the site of the Schiff base linkage with 11-*cis* retinal (Palczewski et al., 2000; Sakmar et al., 2002), and E113, the counter-ion to the Schiff base in the dark state (Sakmar et al., 1989), as well as C110 and C187, which form a critical disulfide bond in the protein (Karnik and Khorana, 1990). Both cone opsin genes also have the conserved P189 residue which is critical for faster cone opsin pigment regeneration (Kuwayama et al., 2002).

Interestingly, *P. melanoleucus* RH1 has serine at site 185 instead of the highly conserved cysteine, similar to several other snakes (Schott et al., 2016; Simões et al., 2016). Mutations at site 185 have been shown to reduce both visual pigment stability (McKibbin et al., 2007) and transducin activation *in vitro* (Karnik et al., 1988). Also, the *P. melanoleucus* RH1 has N83 and S292, which are often found in rhodopsins with blue-shifted λ_max_ values, and can also affect all-trans retinal release kinetics following photoactivation (Bickelmann et al., 2012; van Hazel et al., 2016).

Based on known spectral tuning sites in LWS, *P. melanoleucus* has A285, compared to T285 in *Thamnophis* snakes. T285A is known to blue-shift the LWS pigment by 16-20 nm (Asenjo et al., 1994; Yokoyama, 2000). This suggests that the *P. melanoleucus* LWS may be considerably blue-shifted relative to the LWS pigment in *Thamnophis* snakes. Within *P. melanoleucus* SWS1, the phenylalanine at site 86 suggests that the pigment will be absorbing in the UV, as is typical of reptilian SWS1 pigments (Hauser et al., 2014). *Pituophis melanoleucus* SWS1, as well as other colubrids SWS1 (Simões et al., 2016), have hydrophobic residues at two spectral tuning sites, A90 and V93. These sites are usually have polar or charged amino acid side chains (Carvalho et al., 2011; Hauser et al., 2014). These functional significance of these hydrophobic residues have yet to be characterized, and suggests that caution should be taken in applying spectral tuning predictions on squamates SWS1 pigments.

### Immunohistochemistry

Because *P. melanoleucus* has an all-cone retina, we used immunohistochemistry to determine if both rhodopsin and the rod G protein transducin are expressed in cone photoreceptors. We performed fluorescent immunohistochemistry on the transverse cryosections of the retina of *P. melanoleucus* with the rhodopsin antibody (RET-P1) and a rod-specific transducin antibody (K20). Both antibodies have been previously shown to be selective across a range of vertebrates (Fekete and Barnstable, 1983; Hicks and Barnstable, 1987; Osborne et al., 1999; Schott et al., 2016). We also used these antibodies on a CD1 mouse retina, following similar preparation, as a positive control.

Our results showed rhodopsin localized to the outer segments of select photoreceptors of the *P. melanoleucus* retina (red, Fig 1D), whereas the rod transducin localized to the inner segment (green, Fig 1E). The small amount of colocalization between rhodopsin and transducin in the inner segment (yellow, Fig 1F) is expected as the animal wasn’t dark-adapted prior to sacrifice, as rod transducin translocates to the inner segment when exposed to bright light(Calvert et al., 2006; Elias et al., 2004). This pattern is consistent with rhodopsin and transducin staining in the *T. proximus* retina (Schott et al., 2016) and the previously unexplained results of rhodopsin detected in the retina of *T. sirtalis* (Sillman et al., 1997).

**Figure 1:**
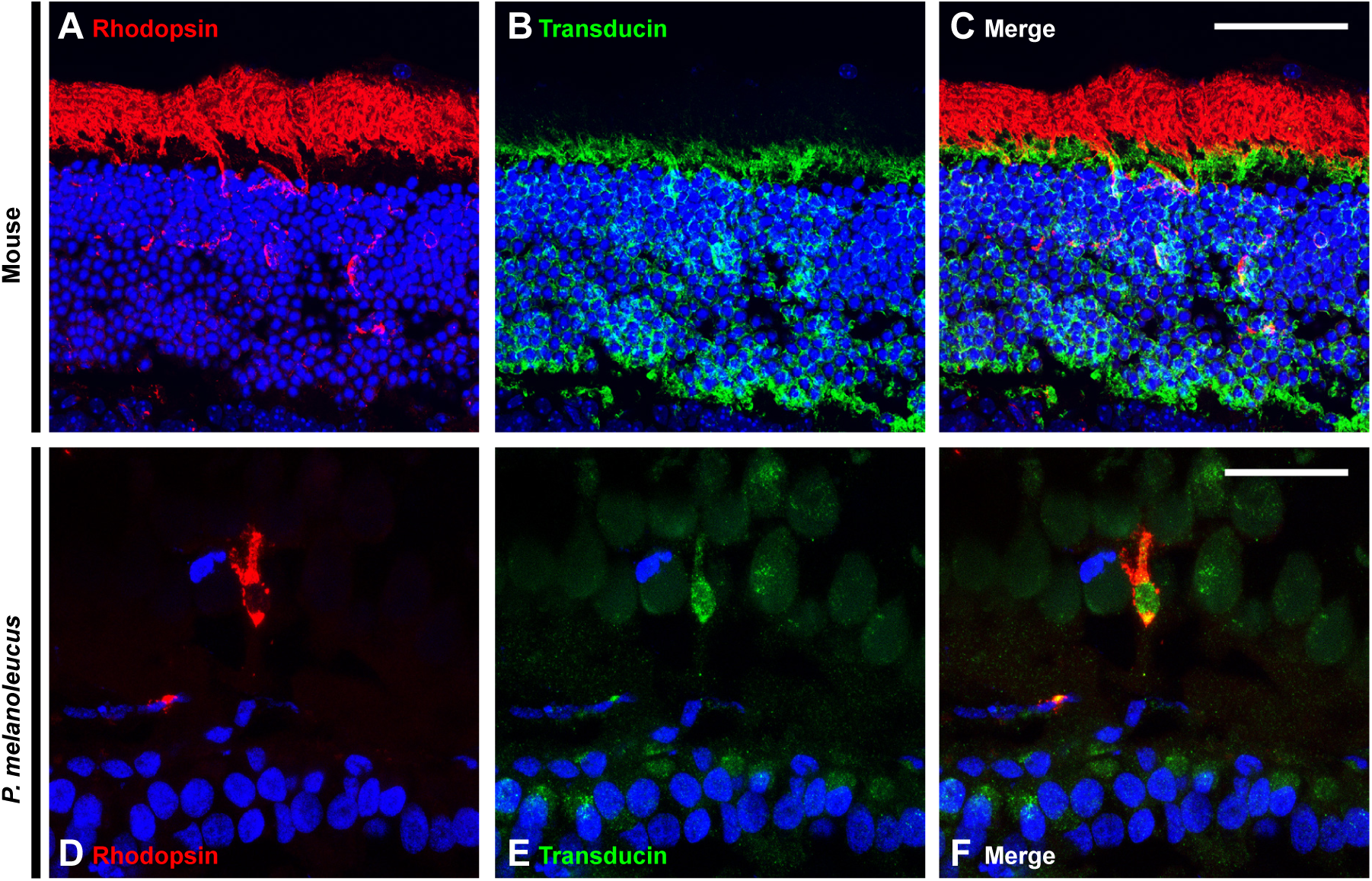
Immunohistochemical staining of control (mouse, **A-C**) and pine snake (**D-F**) transverse retinal cryosections with rhodopsin (RET-P1) and rod-specific-transducin (K20) antibodies. Rhodopsin is found in a subset of cone-like photoreceptors localized to the outer segment (**D**). Rod-specific transducin is also found in the same photoreceptor, primarily to the inner segment (**E**). Double staining indicates that both rhodopsin and rod-specific transducin are found within the same cell (**F**). Nuclei are shown in blue, rhodopsin in red, and rod-specific transducin in green. Scale bars = 20 μm.

As expected, CD1 mouse retina had strong rhodopsin fluorescence (red, Fig 1A) in the outer segment and strong rod transducin staining (green, Fig 1B) in the inner segment, consistent with the rod dominant mouse retina. The lack of colocalization is consistent with a light-adapted retina (Calvert et al., 2006; Elias et al., 2004) (Fig 1C).

### In vitro expression

Complete coding sequences of the *P. melanoleucus* RH1, LWS, and SWS1 opsins were cloned into the p1D4-hrGFP II expression vector (Morrow and Chang, 2010). Expression vectors were then transfected into HEK293T cells and the expressed protein was purified with the 1D4 monoclonal antibody (Morrow and Chang, 2015; Morrow et al., 2011). Bovine wildtype rhodopsin was used as a control (Fig 2A). Pine snake rhodopsin has a λ_max_ of 481nm (Fig 2B) which is similar to the measured λ_max_ of rhodopsins from *T. proximus, T. sirtalis*, and *Arizona elegans* snakes (Schott et al., 2016; Sillman et al., 1997; Simões et al., 2016). The drastic blue shift is expected given the presence of the blue-shifting N83 and S292 amino acid identities (Bickelmann et al., 2012; Dungan et al., 2016; van Hazel et al., 2016). *P. melanoleucus* rhodopsin expressed similar to that of *T. proximus*, with a large ratio between total purified protein (absorbance at 280nm) and active protein (absorbance at λ_max_) that indicates that only a small proportion of the translated opsin protein is able to bind chromophore and become functionally active. One possible explanation for this effect is the S185 residue in *P. melanoleucus* rhodopsin, as mutations at this site have been shown to affect the retinal binding efficiency of rhodopsin pigments expressed *in vitro* (McKibbin et al., 2007).

**Figure 2:**
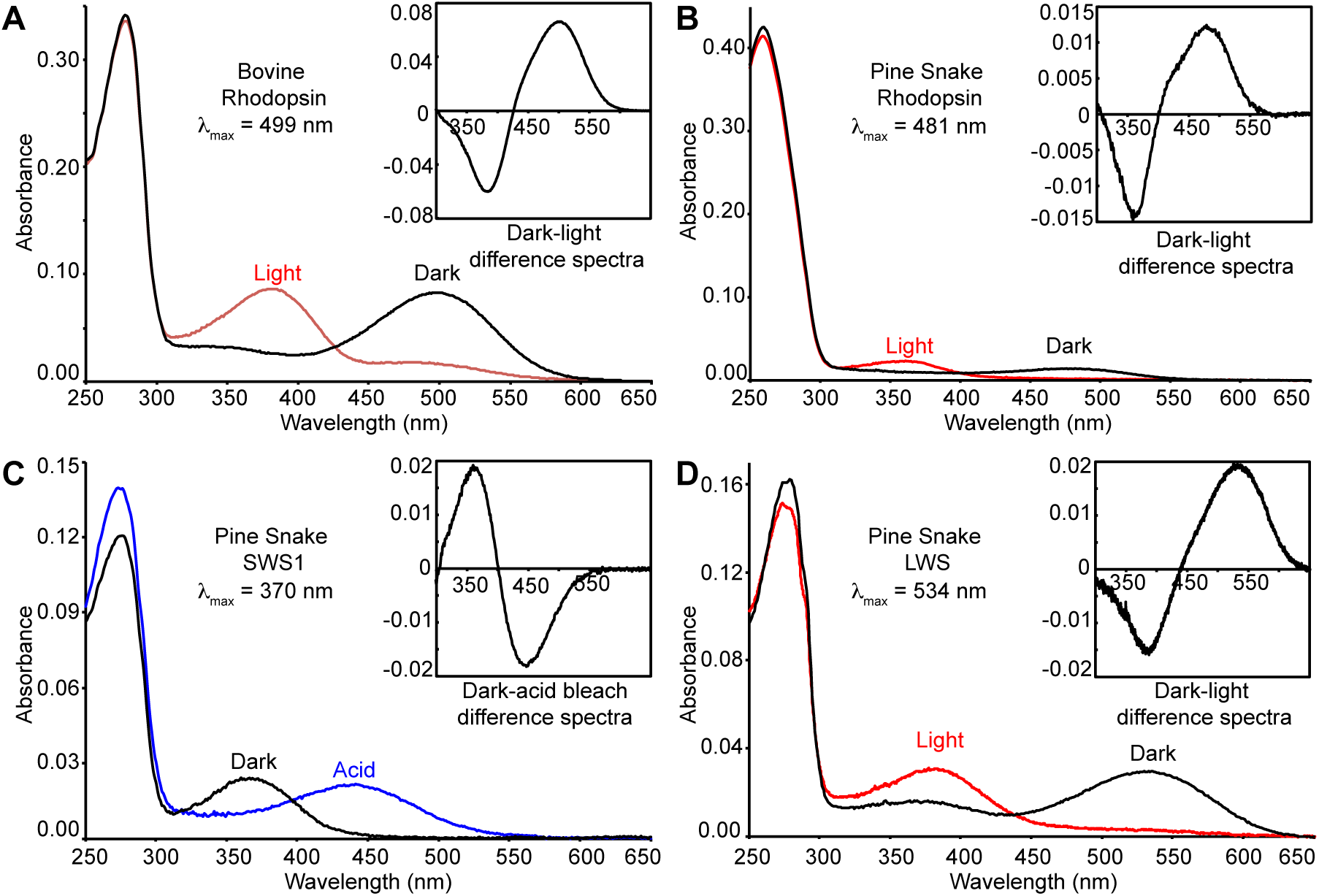
UV-visible dark absorption spectra of pine snake opsins. (**A**) Bovine wildtype rhodopsin was used as a control for expressions. Dark spectra for pine snake (**B**) rhodopsin (**C**) SWS1 and (**D**) LWS. Inset in (**A**), (**B**), and (**D**) is the dark-light spectra. Inset in (**C)** is the dark-acid bleach spectrum. λ_max_ estimations are shown for each pigment.

Expression of pine snake SWS1 showed a much more favorable 280nm to λ_max_ ratio (Fig 2C). We found that *P. melanoleucus* SWS1 pigment absorbs in the UV range with a λ_max_ of 370nm, similar to the SWS1 λ_max_ of *Lampropeltis getula, Rhinocheilus lecontei,* and *Hypsiglena torquata* (Simões et al., 2016) all of which have the most red-shifted UV SWS measured among colubrid snakes.

Similar to the SWS1 expression, LWS also expressed quite well (Fig 2D). Fit to A1 templates gave a λ_max_ of 534nm, which is blue-shifted relative to *Thamnophis* (Schott et al., 2016; Sillman et al., 1997), but identical with LWS MSP measurements of *H. torquata* (Simões et al., 2016) and very close to those of *L. getula, A. elegans*, and *R. lecontei* (Simões et al., 2016).

### Opsin protein functional characterization

In order to confirm the covalent attachment of the chromophore in *P. melanoleucus* SWS1 pigments, the purified opsin was acid bleached (Fig 2C). We found a shift of the λ_max_ from 370nm to 440nm, which indicates the presence of 11-*cis* retinal covalently bound by a protonated Schiff base to a denatured opsin protein (Kito et al., 1968), suggesting that the UV sensitivity of the pigment may be established by only the presence of F86.

*P. melanoleucus* LWS (Fig 3A) and RH1 (Fig 3B) were tested for hydroxylamine reactivity, which assesses the accessibility of the chromophore-binding pocket to attack by small molecules. If hydroxylamine can enter the binding pocket, it will hydrolyze the Schiff base linkage, resulting in an absorbance decrease of the dark peak and the increase of a retinal oxime peak at 363nm. Rhodopsins are thought to be largely non-reactive in the presence of hydroxylamine (Dartnall, 1968) (Fig 3C) due to their highly structured and tightly packed chromophore binding pockets relative to cone opsins, which often react when incubated in hydroxylamine (van Hazel et al., 2013). *P. melanoleucus* LWS reacted to hydroxylamine, as expected, with a t_1/2_ of ~3.9 min (Fig 3A), a time within the range of cone opsins (Das et al., 2004; Ma et al., 2001). As the λ_max_ of *P. melanoleucus* SWS1 is 370nm, it was not tested as we would not be able to distinguish the retinal oxime peak from the λ_max_ peak. Interestingly, *P. melanoleucus* rhodopsin also reacted to hydroxylamine with a t_1/2_ of ~14 min (Fig 3B), unlike the bovine rhodopsin control that did not react (Fig 3C). This implies that the chromophore binding pocket of *P. melanoleucus* rhodopsin has a more open configuration relative to other rhodopsin proteins, a property more typical of cone opsins.

**Figure 3:**
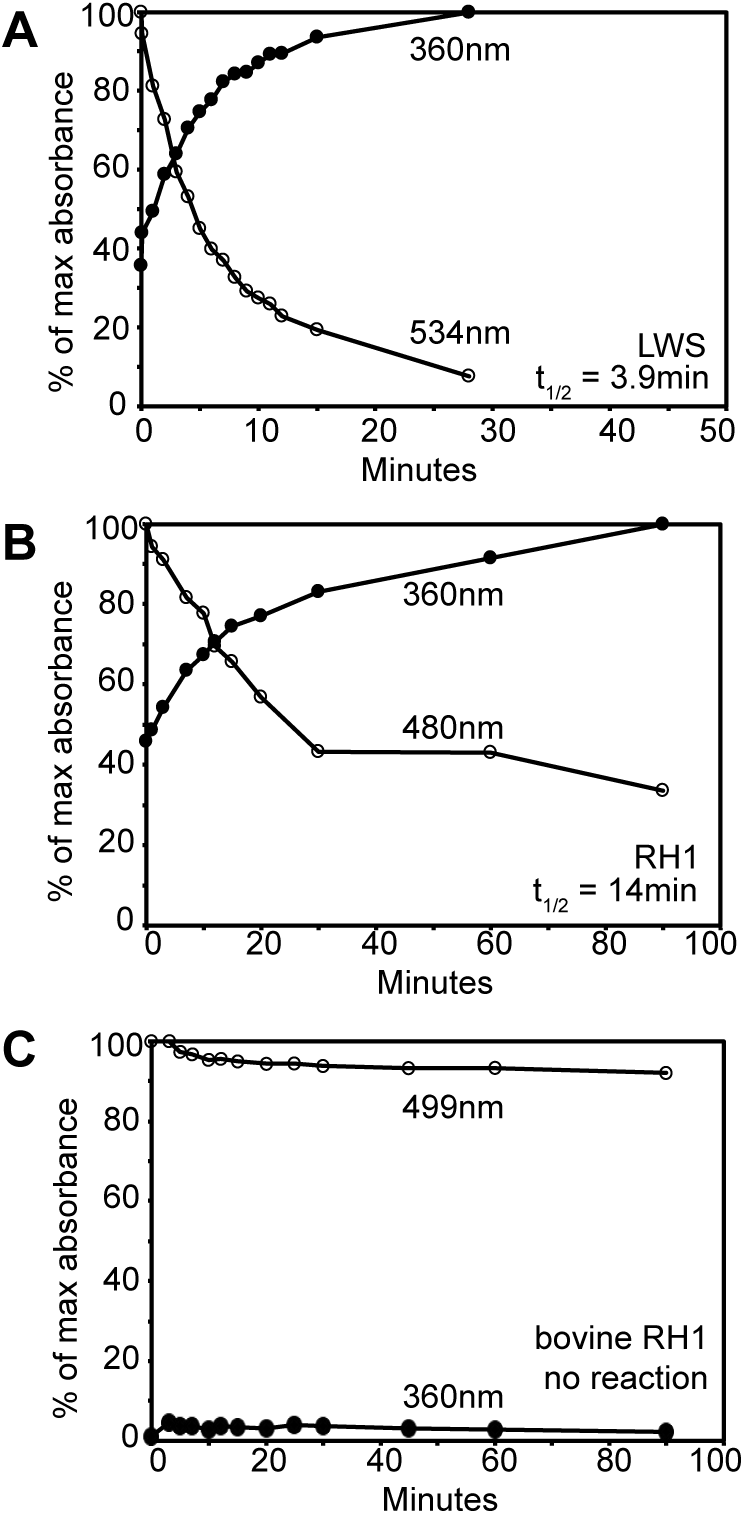
Hydroxylamine reactivity of pine snake (**A**) LWS and (**B**) RH1 pigments and (**C**) bovine rhodopsin. Absorption values of the dark λ_max_ peak decrease over time (open circles), while absorption of the retinal oxime at 360nm increase over time (solid circles). The half-lives of the reactive opsins were determined via curve fitting exponential rise and decay equations to data.

## Discussion

Recently, there has been mounting evidence supporting the theory of transmutation in photoreceptor evolution, proposed by Walls in 1942, which outlines the evolutionary transformation of one photoreceptor type into another in reptilian retinas. Evidence of cone to rod transmutation in nocturnal geckos has been extensively demonstrated using both cellular and molecular techniques (Crescitelli, 1956; Dodt and Walther, 1958; Kojima et al., 1992; McDevitt et al., 1993; Röll, 2001; Sakami et al., 2014; Tansley, 1959; Tansley, 1961; Tansley, 1964; Zhang et al., 2006), while evidence of rod-to-cone transmutation in colubrid snakes remains somewhat sparse (Schott et al., 2016). In order to demonstrate rod-to-cone transmutation in the retina, there needs to be evidence of a functional rod machinery in a photoreceptor with some rod-like features in a retina with appears, superficially, to consist of only cones. Certainly, the presence of RH1 genes and MSP data suggests transmutation has occurred in several colubrid species (Hart et al., 2012; Sillman et al., 1997; Simões et al., 2015; Simões et al., 2016), but further investigation is required in order to firmly state transmutation is present in the retinas of these colubrid snakes as there are multiple alternate explanations possible (RH1 in the genome but not expressed, rhodopsin expressed but not functional, a cone cell co-opting rhodopsin etc). There is only one colubrid snake species for which cellular and molecular evidence for transmutation has been reported, *Thamnophis
proximus* (Schott et al., 2016).

This study provides strong evidence that supports the hypothesis that photoreceptor transmutation has occurred in the retina of *P. melanoleucus*. As *P. melanoleucus* is not closely related to snakes in the genus *Thamnophis*, this suggests that transmutation may be widespread in colubrid snakes. However, the functional significance of transmutation in colubrid snakes still has not been established. In geckos, the advantage of cone-to-rod transmutation is more straightforward as these nocturnal animals are most likely compensating for the loss of RH1 in their diurnal ancestor. We propose that transmutation in colubrids may have occurred as an adaptation to diurnality that provided *P. melanoleucus* with a cone-like rod photoreceptor that operates at brighter light levels, perhaps as a compensation for the loss of the RH2 cone opsins. Our finding of a highly blue-shifted rhodopsin with more cone-like functional properties, as indicated by hydroxylamine reactivity, support this hypothesis.

*Pituophis melanoleucus* rhodopsin shows hydroxylamine reactivity, a canonical cone opsin property (Wald et al., 1955). With a reaction half-life of ~14min, the *P. melanoleucus* rhodopsin reacts much quicker and closer to cone opsin speeds (Das et al., 2004; Ma et al., 2001) than previous rhodopsins that have reacted when incubated in hydroxylamine, like the echidna (Bickelmann et al., 2012) and the anole (Kawamura and Yokoyama, 1998) which react over hours. The RH1 sequence contains both E122 and I189, which are known to mediate the slower decay and regeneration kinetics typical of rhodopsin (Imai et al., 1997; Kuwayama et al., 2002). Conversely, the presence of serine rather than cysteine at site 185, in rhodopsin has been shown to activate fewer G proteins (Karnik et al., 1988) and mutation at site 185 has been shown to reduce the thermal stability of the protein (McKibbin et al., 2007), both characteristics being more typical of cone opsins. Cones have been optimized for fast regeneration, with cone opsin meta-intermediate states being short lived compared to rhodopsin (Imai et al., 2005), and a cone-specific Müller cell retinoid cycle (Das et al., 1992) providing a dedicated pool of 11-*cis* retinal. These faster kinetic properties are hypothesized to be facilitated in cone opsins via the relative “openness” of the chromophore binding pocket, which allows water molecules, and therefore other small molecules like hydroxylamine, to access the chromophore where they can participate Schiff base hydrolysis (Chen et al., 2012; Piechnick et al., 2012; Wald et al., 1955). Rhodopsins, on the other hand, are optimized for sensitivity and signal amplification; therefore, E122/I189 and a tighter overall structure contribute to a slower active state decay allowing for the activation of multiple G proteins (Chen et al., 2012; Starace and Knox, 1997), increased thermal stability relative to cone opsins (Barlow, 1964), and a resistance to hydroxylamine (Dartnall, 1968). *Pituophis melanoleucus* rhodopsin shows adaptations to decrease the number of G-proteins activated, and hydroxylamine reactivity which suggests that an open chromophore binding pocket would enable water access to facilitate active state decay, Schiff base linkage hydrolysis, and retinal regeneration (Chen et al., 2012) rates similar to cone opsins. Spectroscopic assays measuring G protein activation and retinal release rates have never been performed on colubrid rhodopsins, but would be an interesting direction for future research characterizing this cone opsin-like rhodopsin.

Retinal immunohistochemistry localized *P. melanoleucus* rhodopsin protein in the outer segment of a photoreceptor, as well as the presence of rod transducin in the inner segment. Rod and cone transducin are thought to originate via duplication from on ancestral gene (Larhammar et al., 2009) and both have been shown to function with all opsins (Sakurai et al., 2007), therefor the presence and preservation of rod transducin in the photoreceptor supports the theory that this is indeed a transmuted rod and not a cone photoreceptor co-opting rhodopsin expression. Because the retinas were not dark adapted prior to sacrifice, we can presume that under normal photopic light conditions, *P. melanoleucus* rod transducin is cycled out of the outer segment of the cone-like rod, a distinct rod property (Chen et al., 2007; Rosenzweig et al., 2007). In the light, rods cycle transducin and recoverin out of the outer segment, and arrestin into it (Calvert et al., 2006). This allows the rod to effectively shut down phototransduction under bleaching conditions to prevent damage to the photoreceptor. Cones generally do not cycle transducin out of the outer segment of the photoreceptor under normal light conditions (Chen et al., 2007). This suggests that the rhodopsin-expressing photoreceptors in the retina of *P. melanoleucus* would not be able to generate a photoresponse in normal daylight, and thus if this cone-like rod is participating in colour vision with the canonical cones in the retina, it would likely only be under mesopic light conditions where both photoreceptor cell types can be active.

Our microscopy results of the *P. melanoleucus* retina additionally revealed a cone-like rod which still looks distinct in comparison to the other cones. The cone-like rod outer segment and inner segment had similar diameters with a relatively long outer segment, while the surrounding cones had distinctly large ellipsoids in the inner segment, and proportionally smaller outer segments. Rod photoreceptor morphology is also generally specialized to maximize sensitivity with long cylindrical outer segments (Lamb, 2013). Cone morphological specializations, however, are thought to enable selective colour vision, a faster phototransduction and visual pigment regeneration, while also minimizing metabolic load by miniaturizing the overall structures with large ellipsoids that tunnel light onto smaller tapered outer segments (Harosi and Novales Flamarique, 2012). Previous EM studies on the retina of *T. proximus* showed that the membrane discs unique to rods in the outer segment are still present in the transmuted photoreceptor (Schott et al., 2016). Interestingly, a reduction of RH1 expression levels has been shown to reduce the size of the outer segment of rods, in addition to lowering the photosensitivity and altering the kinetics of the cell to be more cone-like (Makino et al., 2012; Rakshit and Park, 2015; Wen et al., 2009). Currently, the relative expression levels of RH1 in the retinas of colubrid snakes have not been measured. There are additional specializations in the synaptic structures that reflect the different priorities in rod and cone function (Lamb, 2013), but the synaptic structure of the cone-like rod also remains uninvestigated.

Results from this study suggest that transmutation is modifying the function of a subset of photoreceptors in the retina of *P. melanoleucus*. These modifications may serve to lower the sensitivity and signal amplification of the photoreceptor, supporting the hypothesis of a more cone-like function. However, the type of signal these transmuted rods send to the brain is still unknown. Rods and cones are known to have distinct ERG responses, but *T. sirtalis* is the only colubrid snake with ERG measurements performed at a variety of light levels (Jacobs et al., 1992). However, this study did not record any scotopic (rod) response, nor did it record any photopic response from the SWS1-type photoreceptors, which suggests that the results of the study may be incomplete or that the scotopic pathways in the colubrid eye have degraded. Indeed, in high scotopic and mesopic light levels, mammalian rod photoreceptors can and do use cone pathways (Kolb et al.). The presence of rod bipolar cells and AII amacrine cells, both of which are required in the rod-specific photoresponse pathway (Lamb, 2013), has never been established in the colubrid retina.

Transmutation may be an attempt to compensate for the loss of the RH2 cone opsin and the lack of spectral overlap between the LWS and SWS1 pigment, such that the rod photoreceptor may have evolved cone-like functionality such that it could participate in colour vision. In addition to the molecular modifications to *P. melanoleucus* rhodopsin and the physiological modifications to the rod cell, the extreme blue-shift of the RH1 λ_max_, which is quite rare for terrestrial rhodopsins, may itself be an adaptation for colour vision, as a λ_max_ of ~480 nm is in the range of typical RH2 pigments (Lamb, 2013). *Pituophis melanoleucus*, in comparison to the *Thamnophis* genera (Schott et al., 2016; Sillman et al., 1997), has a narrower overall range of spectral sensitivities. There could be two possible reasons for this narrowing. It could be that this narrowing of the spectral ranges is to facilitate spectral overlap as an adaptation in *P. melanoleucus*. Or the narrowing of the spectral range may simply be due to phylogenetic history, as *P. melanoleucus* LWS and SWS1 absorb at similar wavelengths to its closest relatives (Simões et al., 2016), which in turn could be an adaptation, but not one due to the specific visual environment of *P. melanoleucus*. Trichromatic vision would be greatly advantageous for a diurnal species (Ankel-Simons and Rasmussen, 2008; Heesy and Ross, 2001), and perhaps sacrificing scotopic vision in order to achieve better mesopic and photopic vision is possible, since other snake sensory systems adaptations, such as chemoreception, could be sufficient in dim light environments (Drummond, 1985). However, currently there is a lack of behavioral studies investigating trichromatic colour discrimination in colubrid snakes under mesopic light conditions.

We hypothesize that rod-to-cone transmutation may be allowing colubrid snakes to have a third cone-like photoreceptor, allowing for spectral sensitivity between SWS1 and LWS, possibly also trichromatic colour perception in mesopic light conditions. The loss of RH1 in nocturnal geckos and the resulting transmutation of cone into rod demonstrate that the visual system of squamates is capable of adapting to compensate for previous functionality loss in different photoreceptor types. In colubrid snakes, and possibly squamates in general, the rod/cone photoreceptor binary is not as distinct as it is in other vertebrates and caution should be taken in classifying rod or cone photoreceptors based on limited characterization.

In summary, we find that *P. melanoleucus*, like *T. proximus*, has an all-cone retina derived through evolutionary transmutation of the rod photoreceptors. Furthermore, *P. melanoleucus* rhodopsin is the first vertebrate rhodopsin to show hydroxylamine reactivity similar to cone opsins. This study is also the first to demonstrate the functional effects of transmutation in the retina of colubrid snakes. We suggest that transmutation in colubrid snakes is an adaptation to diurnality and is compensating for the loss of RH2 by establishing spectral sensitivity in a range where the existing SWS1 and LWS are not sensitive, and possibly establishing trichromatic colour vision. Perhaps transmutation in colubrid snakes may have contributed to the widespread success of the snake family across such a vast range of ecologies and lifestyle. Ultimately, future work investigating the functional effects of transmutation, from behavioral to molecular, will reveal the significance of rod-to-cone transmutation in colubrid snakes.

## Author contributions

N.B and B.S.W.C contributed to the conceptual design. V.T. provided mice and experimental advice. N.B. and B.D. performed the research, and N.B., R.K.S, and B.S.W.C analyzed data. N.B., R.K.S and B.S.W.C wrote the manuscript.

